# Reduced wound healing and angiogenesis in aged zebrafish and turquoise killifish

**DOI:** 10.1101/2022.08.16.503961

**Authors:** Johanna Örling, Katri Kosonen, Jenna Villman, Martin Reichard, Ilkka Paatero

## Abstract

Impaired wound healing is associated with aging and has significant effects on human health on an individual level, but also the whole health care sector. Deficient angiogenesis appears to be involved in the process, but the underlying biology is still poorly understood. This is at least partially being explained by complexity and costs in using mammalian aging models.

To understand aging-related vascular biology of impaired wound healing, we have utilized zebrafish and turquoise killifish fin regeneration models. The regeneration of caudal fin after resection was significantly reduced in old individuals in both species. Age-related changes in angiogenesis, vascular density and expression levels of angiogenesis biomarker VEGF-A were observed. Furthermore, anti-angiogenic drug, vascular endothelial growth factor receptor blocking inhibitor SU5416 reduced regeneration indicating a key role for angiogenesis in the regeneration of aging caudal fin despite aging-related changes in vasculature.

Taken together, our data indicates that these fish models are suitable for studying aging-related decline in wound healing and associated alterations in aging vasculature.

## Introduction

Aging is a feature affecting most, if not all, contemporary human populations. Aging, however, causes many physiological and biological changes in individual and is also considered as one of the primary risk factors for complicated wound healing ^1^. Challenged wound healing or chronic wounds are sometimes called a silent epidemic and they create a significant burden to affected individuals and also to the health care sector ^2^. In developed countries the treatment expenses due to chronic wounds cover up to 5.5% of the healthcare costs ^3,4^.

Wound healing consists of four overlapping phases called hemostasis, inflammation, proliferation and remodeling, and complications in any of these phases contribute to poor wound healing ^5^. One of the crucial processes during wound healing is angiogenesis. Angiogenesis is required for tissue remodeling, and it initiates rapidly after wounding ^6^. Angiogenic factors, such as VEGF, promote the action of endothelial cells for instance by stimulating their migration and proliferation ^7,8^. Hypoxia plays a significant role in wound healing^9^ and hypoxia inducible factor 1 (HIF-1α) acts as a stimulus for the production of VEGF ^10^. As wound healing enters the final phases, high levels of growth factors start to decrease, hypoxia subsides and angiogenesis is suppressed ^11^. During aging, impaired HIF-1α signaling indicates a diminished response to hypoxia and thus plays a role in the delays concerning angiogenesis ^12^. In addition, increased secretion of proinflammatory mediators due to the dysfunction of macrophages impairs the production of VEGF in elderly ^13^. Interestingly, impaired VEGF signaling has been suggested to play a key role in aging process ^14^.

Although the underlying etiologies behind chronic wounds vary, they share many similarities, such as senescent cells unable to proliferate, dysfunctional stem cells, high levels of proteases and reactive oxygen species, increased levels of proinflammatory cytokines, and lack of microvasculature ^5^. Lack of microvasculature may result from poor angiogenesis, and angiogenesis is compromised during aging although the underlying mechanisms are largely not understood ^15^.

Similarly to angiogenesis ^15^, the mechanisms of wound healing have been studied in great detail in young individuals, but the defects in biological processes resulting in poor wound healing in aged individuals are still unclear in many ways ^16^. One reason for this may be the long lifespan of laboratory mammals, which complicates aging studies. On the other hand, short-lived invertebrates lacking blood vessels may have limitations in potential to translate the findings into mammalian and human systems. In recent years, laboratory fish have been utilized as models for wound healing and tissue regeneration. Most commonly used model fish, zebrafish (*Danio rerio*), has fairly long lifespan but another shorter lived model species, turquoise killifish (*Nothobranchius furzeri*), is gaining ground as suitable vertebrate model for aging studies ^17,18^.

Here, we have utilized zebrafish and turquoise killifish as complementary models to study aging-related decline in wound healing by using fin regeneration model. We have especially focused on the alterations of angiogenic signalling and vasculature associated with compromised wound healing in this study.

## Materials and Methods

### Fish husbandry and ethics

The authorization for this project was obtained from Animal Experimental Board of Regional State Administrative Agency for Southern Finland (ESAVI/16458/2019 and ESAVI/2402/2021). The turquoise killifish in the facility were maintained according to protocol from Polačik et al. ^19^ by the trained personnel and zebrafish according to standard procedures. Killifish were fed twice a day with frozen blood worms and zebrafish with Gemma micro dry food. Water parameters in turquoise killifish system were following: temperature 26°C, salinity 230 ± 30 μS and pH 7.3-7.6.

### Zebrafish caudal fin regeneration assay

Wild-type male zebrafish (AB line) aged 4 months (12 fish) and 4 years (12 fish) were utilized for studying the differences in wound healing between young and aged fish. At the beginning of the study (day 0), the fish were anesthetized with 200 mg/l tricaine methanosulfonate MS222 (Sigma) diluted in the system water. As the fish showed no physical reactions upon touching, the depth of the anesthesia was considered deep enough. The fish was placed on a petri dish lid with the help of tablespoon, and its caudal was spread carefully on top of the lid. After this, the caudal fin was photographed via Zeiss Stemi DV4 stereomicroscope using smart phone camera and an adapter. Finally, a small piece of distal caudal fin was resected with a scalpel, and the fin was photographed once more. Half of the resected piece of fin was stored at −80°C, and the other half was fixed in 10% neutral buffered formalin. After recovering from anesthesia, all the fish were housed individually in a 1-liter tank, which contained 0.004 % methylene blue in system water. The tanks were placed in a warm room (26°C) for the duration of the experiment. The system water containing methylene blue in the tanks was changed to fresh every other day and the fish were fed just before water change. During the study the fish were anesthetized at days 2, 4, 6 and 8 and their caudal fins were photographed under stereo microscope. At the end of the study (day 8) the regenerated part of distal caudal fin was again amputated under anaesthesia and finally stored at −80°C.

### Turquoise killifish caudal fin regeneration assay

Both genders of turquoise killifish (*Nothobranchius furzeri*, strain MZCS-222, Reichard lab, Institute of Vertebrate Biology, Czech Academy of Sciences, Brno, Czech Republic) was used. The experiment had eight fish in four groups; young male (age 5 weeks), young female (age 5 weeks), old male (age 12 weeks), old female (age 12 weeks). First, the fish were anaesthetized with 400mg/l of tricaine. The distal part of the caudal fin was resected under stereomicroscope (Zeiss Stemi DV4). The caudal fin was imaged before and after resection using a cell phone camera attached to stereomicroscope. The experiment was carried out as described above for zebrafish. The regrowth of fins was imaged on days 1, 2, 3, 4, 7 and 8. After the fish were fed with blood worms, the remaining food and debris was removed with a pipette and half of the water was replaced.

SU5416 (Santa Cruz Biotechnologies) or DMSO (Sigma-Aldrich) alone were dissolved in water. The fish were bathed in drug solution, and 50% of the liquid was replenished daily 1 hr after the feeding of the killifish. The fish were imaged at days 0, 2, 4, 6, and 8.

### Zebrafish caudal fin angiogenesis assay

The analysis of angiogenesis in zebrafish caudal fin was carried out largely similarly as wound healing assays. Zebrafish (*roy, mitfa, Tg(fli1:EGFP)^y1^*) of younger adult population (age 16 months, 10 fish) and old adult population (age 36 months, 8 fish) were used in the experiment. The fins were imaged at 3, 5 and 7 dpa (days post-amputation) and the experiment was ended at 7 dpa after imaging. For imaging, the zebrafish were anesthetized, transferred to a petri dish and imaged using Zeiss AxioZoom fluorescence stereomicroscope with total magnification of 16x. The microscope was equipped with 1.0x PlanApo Z objective, HXP 200C mercury lamp, Alexa 488 - filter set 38 HE, (Ex: BP 470/40nm, Em.: BP 525/50nm) and Hamamatsu sCMOS Orca Flash4.0 LT+ camera. Reflected light images were captured by illuminating the samples with CL 9000 LED CAN ring light against black background and collecting the reflected light without a filter cube.

### RNA extraction and quantitative PCR

The RNA form resected fin pieces was extracted using NucleoSpin RNA Plus XS kit for RNA purification (Macherey-Nagel) according manufacturers protocol. The extracted RNA was stored at −80°C until subjected to reverse transcription using High-Capacity cDNA Reverse Transcription Kit (Applied Biosystems). The obtained cDNA samples were stored at −20 °C until analysed using qPCR.

Primer design for housekeeping gene *tbp* and selected angiogenesis-related genes *cd34* and *vegfa* was carried out using NCBI primer design tool (Ye et al., 2012). Primer sequences were: *tbp(5’-AGCGTTTTGCTGCCGTCATA-3’, 5’-TTGACTGCTCCTCACTTTTGG-3’), cd34(5’-AGATGTGTGCTGCAAGGGCA-3’, 5’-CCTGTAACGTCATCTTCCACG-3’*), and *vegfa(5’-AAGCCGAGAAGATGAGAGCG-3’, 5’-TGTCAGACCAAGGGGAATGC-3’*). Prior to PCR measurements, the cDNA samples were diluted to nuclease-free sterile water. Four replicate PCR reactions from each sample were prepared using PowerUp™ SYBR™ Green Master Mix (Thermo Fisher Scientific), and sterile water was used as a negative control. All the reactions were performed on 96-well plates and analysed using QuantStudio 12kFlex instrument (Applied Biosystems). Raw data was further analysed with Relative Quantification application in Thermo Fisher Cloud service (https://apps.thermofisher.com).

### Histology

The caudal fin pieces fixed in 10% neutral buffered formalin were dehydrated through an ascending series of alcohol, defatted and cleared in xylene and finally infiltrated and embedded to paraffin on a transverse plane. From the paraffin embedded samples, 4 μm sections were cut with the help of a microtome and the sections were collected to microscope slides. Before each staining, xylene was used for removing the paraffin from the sections, and the sections were also rehydrated in a descending series of alcohol. Samples were stained with hematoxylin and eosin according to standard protocol and mounted with Depex (Merck). Samples were imaged using Pannoramic P1000 slide scanner (3DHistech).

### Immunofluorescence stainings

The 4 μm sections cut form the paraffin embedded caudal fin pieces were double stained with anti-phospho-histone H3 (pH3) and collagen-binding adhesion protein 35 (CNA35). To stain and detect the mitotic cells in the tissue, polyclonal rabbit phospho-histone 3 (pH3, ser10) antibody (Cell Signaling) together with Alexa Fluor 647 conjugated anti-rabbit secondary antibody (Invitrogen) were used. To stain collagen, CNA35 (Macherey-Nagel) integrated to pET28a mCherry vector was used. pET28a-mCherry-CNA35 was a gift from Maarten Merkx (Addgene plasmid # 61607)^20^. Molecular weight of the purified CNA35 was 60 kDa and the protein concentration was 7.65 mg/ml. The concentrations used in the staining were 1:200 for pH3, 1:1000 for secondary antibody and 1:100 for CNA35. For dilutions 10% FBS (Gibco) in PBS was used. Following deparaffinization and rehydration of the sections, antigen retrieval was carried out in Retriever 2100 (Aptum Bio) using Universal Buffer (10x, R Universal, Aptum Bio) diluted to the concentration of 1:10 with MilliQ water. Unspecific binding of the antibody in the sections was blocked by incubating the slides in 10% FBS for one hour. Overnight incubation with primary antibody was done at +4 °C and protected from light. Following this, the slides were washed with PBS and incubated with secondary antibody for one hour protected from light at room temperature. DAPI diluted with PBS to the concentration of 0.002 mg/ml was used as a background stain. The slides were incubated with DAPI for 10 minutes in dark at room temperature. Finally, the slides were washed with PBS and mounted with water based Mowiol 4-88 (Calbiochem) mounting media, which also contained 2.5% DABCO (Sigma-Aldrich) as an antifading chemical.

Zeiss AxioZoom V16 fluorescence stereomicroscope with total magnification of 16x was used. The microscope was equipped with 1.0x PlanApo Z objective, HXP 200C mercury lamp, DAPI - filter set 49 (Excitation: G365nm, Emission: BP 445/50nm), Alexa 488 - filter set 38 HE, (Excitation: BP 470/40nm, Emission: BP 525/50nm), Alexa 568 - filter set 45 (Excitation: BP 560/40nm, Emission: BP 630/75nm), Alexa 647 - filter set 50 (Excitation: BP 640/30nm, Emission: 690/50nm) and Hamamatsu sCMOS Orca Flash4.0 LT + camera.

### Image processing

The image analysis and processing were carried out using free image analysis platforms FIJI (Schindelin et al., 2012) and QuPath (Bankhead et al., 2017). QuPath was used for calculating blood vessels in the tissue sections. Vascular density was analysed along the central part of the section along dorsoventral axis, and the correct anterior-posterior location was determined by observed fin ray structures. Immunofluorescence analyses were carried out in FIJI, and collagen area and mitotic cell area were calculated and measured using automatic thresholding. Also, the lengths of the regenerated caudal fins both in zebrafish and turquoise killifish studies were calculated in FIJI. The calculations were done both from the dorsal and ventral side of the fin. Some images were uploaded and handled on OMERO server ^21^. Publication images were produced using OMERO.figure and Inkscape. Images were linearly adjusted for brightness and contrast.

### Statistical analyses

Statistical analyses were carried out using GraphPad Prism (versions 6 and 9). Time-series were analysed using 2-way ANOVA and two-sample comparisons using non-parametric Mann-Whitney test. Correlation between fin ray and blood vessel branch points was analysed using linear regression. Prior to the experimentation, the appropriate sample size for caudal fin regeneration assay was obtained from power calculations using an online calculator (http://powerandsamplesize.com/) using 2-groups, beta (power) 70% or 80% and alfa 5%. Estimated effect size in calculations was 50% and standard deviation 40% resulting in sample sizes of 8 (70% power) to 11 (80% power). Samples were neither formally randomized nor blinded during experimentation. Data was plotted as box-plots indicating median and 1^st^ and 3^rd^ quartiles, whiskers extending from minimum to maximum.

## Results

### Declined regeneration of caudal fin in aged zebrafish

To gain insights into age-associated decline in wound healing we raised zebrafish to very old age (40 months). Then, we carried out zebrafish caudal fin regeneration assay using both of old (40 months) and young (4 months) zebrafish (Fig 1A). Interestingly, the old fish had clearly diminished capability to regenerate resected caudal fins. Both the total regeneration of caudal fin (Fig. 1B) and the thickness of hypopigmented regenerating area (Fig. 1C) were reduced in aged individuals. The regeneration of both dorsal and ventral part of the caudal fin were similarly reduced in aged individuals (Fig. 1D) indicating proper morphological control of regeneration.

**Figure 1.**
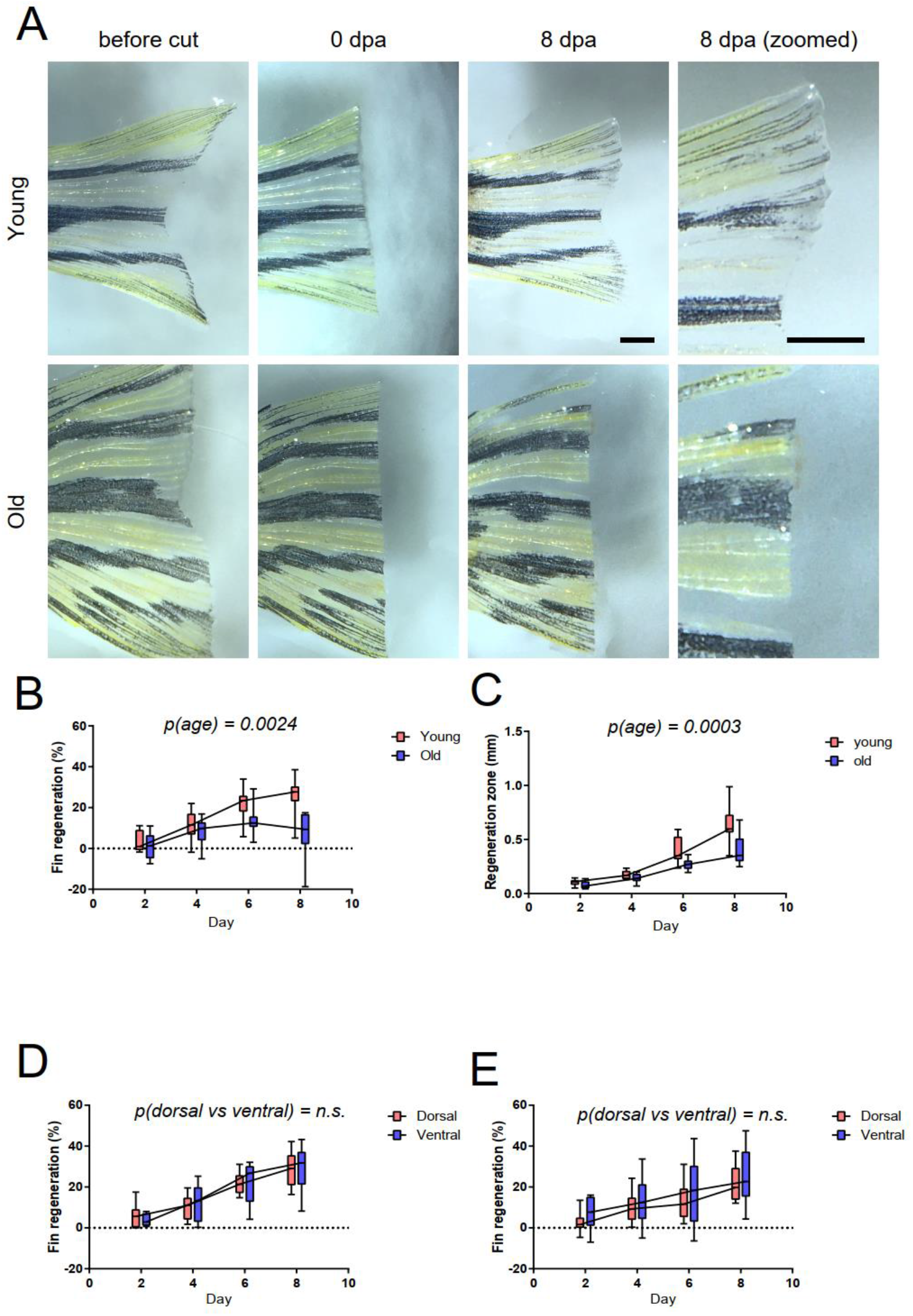
Aging reduces the regeneration of zebrafish caudal fin. Old and young male zebrafish were subjected to caudal fin regeneration assay. A) Microscopy images of fins before, right after and 8 days after resection. Scale bar 1mm. B) Quantification of the total regeneration of the caudal fin. Two-way ANOVA with age and time as factors was used for statistical analysis. C) Quantitation of thickness of regenerating area. Two-way ANOVA with age and time as factors was used for statistical analysis. D, E) Comparison of the regeneration in dorsal and ventral part of caudal fin. Data from young fish in panel D and in old fish in E. n=12 in both age groups. Two-way ANOVA, with age and anatomical location as factors, was used for statistical analysis.

### Declined regeneration of caudal fin in aged turquoise killifish

Zebrafish is a considerably long-lived small fish species and obtaining adequately old zebrafish is a slow process. To overcome these limitations, short-lived turquoise killifish (*Nothobranchius furzeri*) has been proposed as a suitable fish model for aging studies ^17,18^. Therefore, we established a turquoise killifish colony of strain MZCS-222 ^22^ in fish facility in Turku, Finland, to carry out caudal fin regeneration experiments in turquoise killifish. The median lifespan of killifish is known to vary between facilities. In Turku facility the MZCS-222 strain of turquoise killifish had median lifespan 34 weeks (Fig. 2B), which is considerably shorter than approximately 160 week median lifespan of zebrafish ^23^. Caudal fin regeneration experiments were carried out in a similar manner as with zebrafish, with young killifish being 5 weeks and old killifish 12 weeks of age (Fig.2A). Indeed, both the total fin regeneration (Fig. 2C) and thickness of regeneration zone (Fig. 2D) of resected caudal fins of killifish was reduced already in 12-week-old animals compared to 5-week-old young adult fish. These observations were consistent with earlier observations made with a longer-lived killifish strain MZM-0703 ^24^.

**Figure 2.**
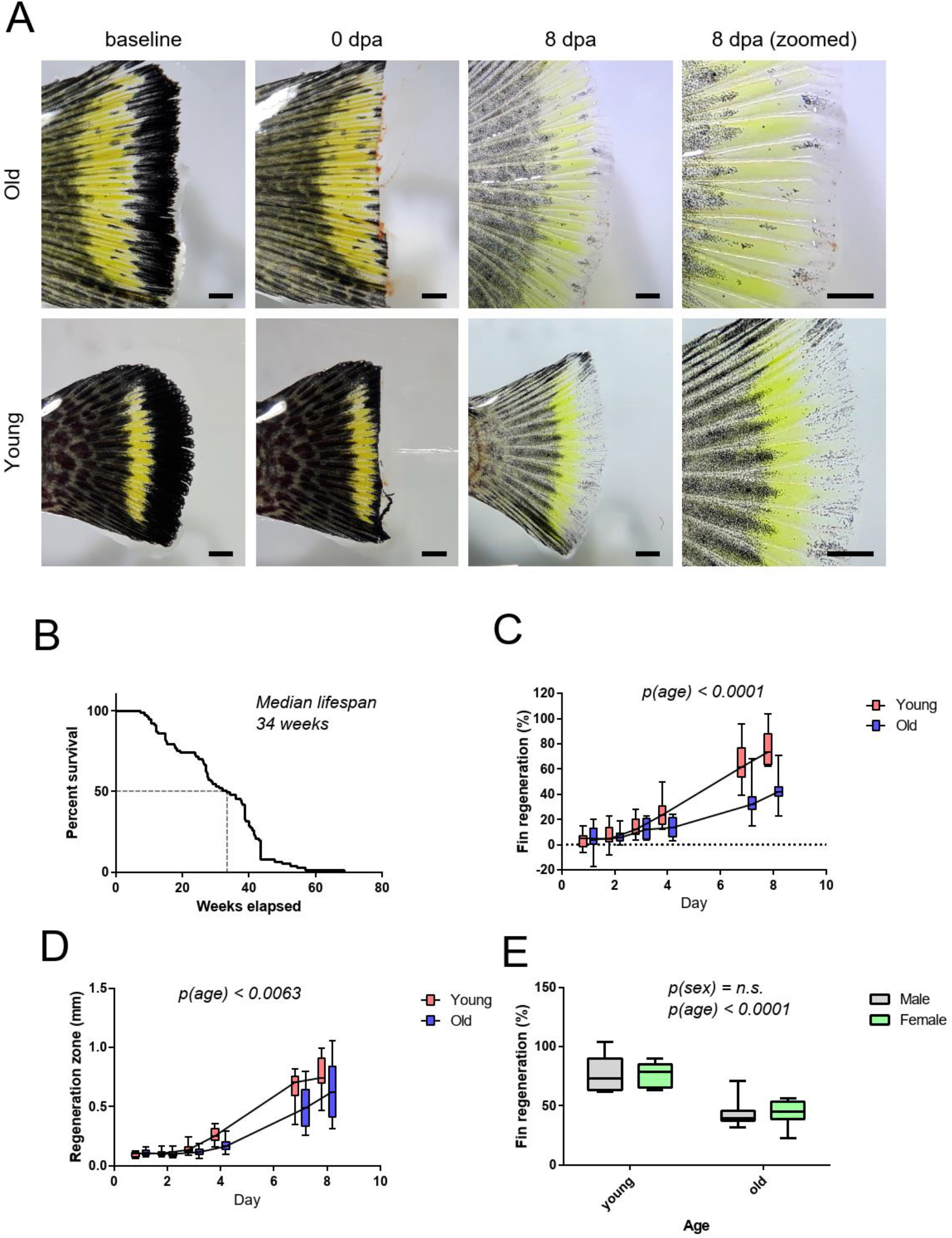
Aging reduces the regeneration of turquoise killifish caudal fin. Old and young turquoise killifish of both sexes were subjected to caudal fin regeneration assay. In total of 4 groups were used (young male, old male, young female, old female). A) Microscopy images of turquoise killifish caudal fins before, right after and 8 days after resection. Scale bar 1mm. B) Kaplan-Meier survival curve of turquoise killifish strain MSCZ222 housed in Turku, Finland. n= 77. C) Quantification of the total regeneration of the caudal fin. Two-way ANOVA with age and time as factors was used for statistical analysis, n=8/group. D) Thickness of regenerating zone. Two-way ANOVA with age and time as factors was used for statistical analysis. E) Comparison of the total regeneration in male and female turquoise killifish caudal fin. In E two-way ANOVA with sex and age as factors was used for statistical analysis. n=8 per group.

There were no differences between the sexes as both males and females displayed similar growth rates and also similar decline of regeneration in old individuals (Fig 2E). To our knowledge, this is the first observation addressing potential sex differences in wound healing in turquoise killifish. In zebrafish, the sex of the animals is not always documented during wound healing and fin regeneration experiments. However, some evidence implies that in general sex differences could be small or absent during wound healing of zebrafish caudal fin, but may also differ in other tissues such as pectoral fins ^25^.

### The vascular density increases in aged individuals

To obtain deeper insights into regeneration process, we analysed tissue sections of fins from old and young killifish. In histological analyses, we surprisingly observed increased blood vessel density in killifish fins during aging (Fig 3A and B). The analysis of vasculature from these tissue sections was, however, quite challenging and we were able to detect only patent blood vessels occupied with red blood cells. To confirm observation of the age related changes in vasculature and angiogenic potential, we analysed old and young transgenic zebrafish carrying EGFP in vascular endothelium (*roy, mitfa, Tg(fli1:EGFP)*) using fluorescence microscopy. This analysis indicated that the vascular density was increased also in zebrafish during aging (Fig 3C and D). The number of branching points was increased in aged blood vessels (Fig. 3E), explaining the increased vascular density. In normal conditions, the caudal fin vasculature is largely organised along fin rays ^26^. Therefore, we analysed if the increased vascular density was actually a result of increased branching of fin rays or from ectopic vascular branches. In the images of fin rays and vasculature (Fig. 3F), we observed similarly increased branching in fin rays (Fig. 3G) which was tightly correlated with increased vascular branching (Fig. 3H). This indicated that increased baseline vascular density is likely not a result of increased angiogenesis and formation of new ectopic vessels, but rather a secondary result of increased branching of fin rays.

**Figure 3.**
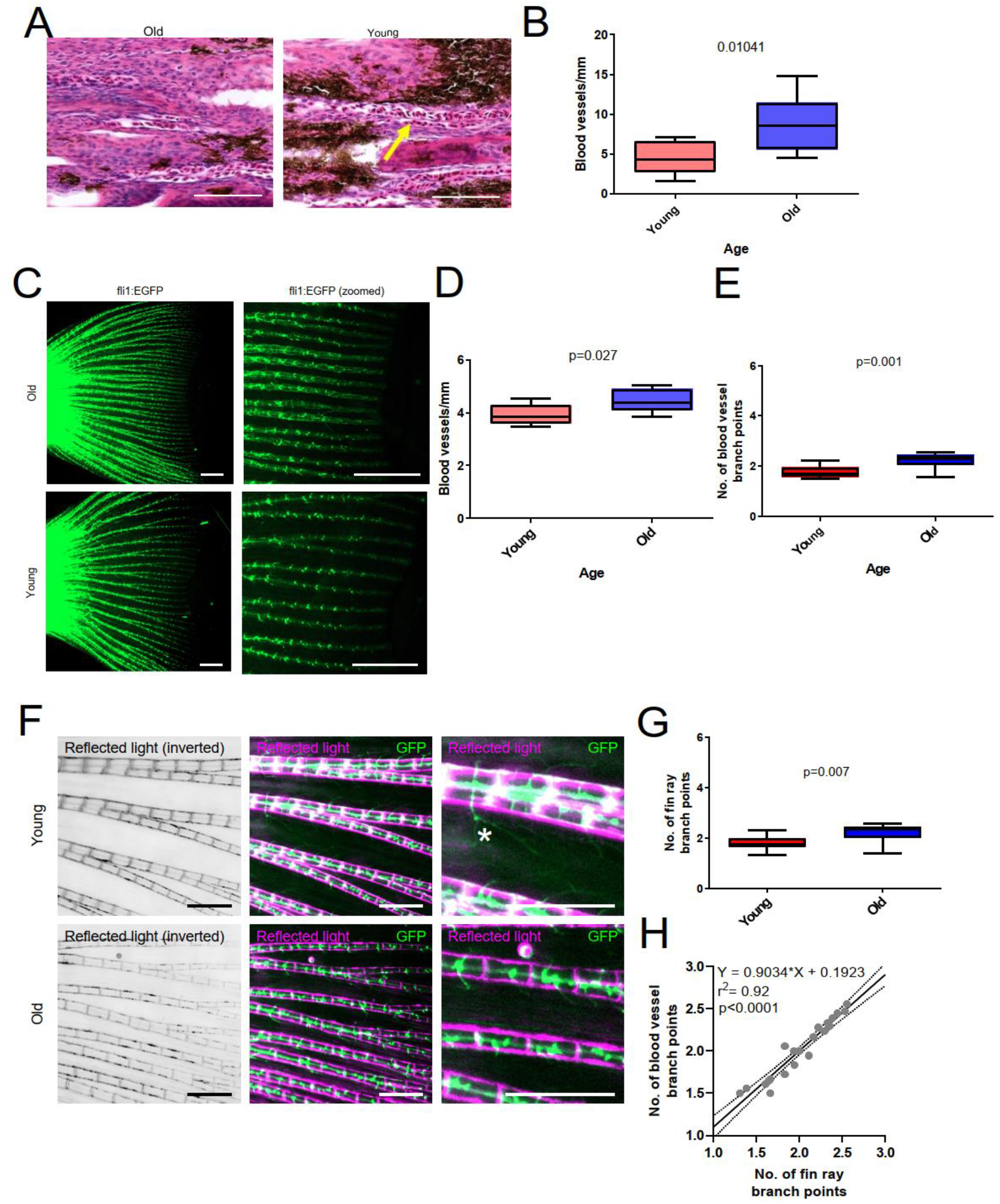
Aging increases vascular density in caudal fins. A) Images of blood vessels in histological section of old and young turquoise killifish caudal fins stained with H&E. An example of a blood vessel is indicated with a yellow arrow. Scale bar 100 μm. B) Quantification of blood vessel density in turquoise killifish sections. Statistical analysis with Mann-Whitney U-test, n=8 per group. C) Fluorescence images of vasculature of transgenic zebrafish (*roy, mitfa, Tg(fli1:EGFP)*) caudal fins. Old fish were 36 weeks (n=8) and young fish 16 weeks (n=10). Scale bar 1mm. D) Quantification of blood vessel density in transgenic zebrafish caudal fins. Statistical analysis with Mann-Whitney U-test. E) Quantification of major blood vessel branch points in transgenic zebrafish caudal fins. F) Fluorescence and reflected white light illumination images of transgenic zebrafish caudal fins. G) Quantification of branching of fin rays. Scale bar 0.5 mm. H) Correlation analysis of fin ray and vascular branches. Statistical analysis using linear regression, both old and young samples combined, n=18.

### Angiogenic potential declines in aged fins

While closely examining the fluorescence microscopy images of young zebrafish (Fig. 3F), we identified some prominently long endothelial protrusions indicative of active angiogenesis. To more directly analyse the capability for angiogenesis, we carried out zebrafish caudal fin angiogenesis assay ^26^ using transgenic zebrafish *roy, mitfa, Tg(fli1:EGFP)*). The distal part of the caudal fin was resected as before, and the regrowth of blood vessels was followed using fluorescence microscopy. Interestingly, the newly developed vascular sprouts during fin regeneration were shorter in aged fish (Fig. 4A and B). This indicated that the angiogenesis was impaired in aged zebrafish. Despite of higher density of blood vessels in aged fins, their angiogenic potential seemed to be diminished.

**Figure 4.**
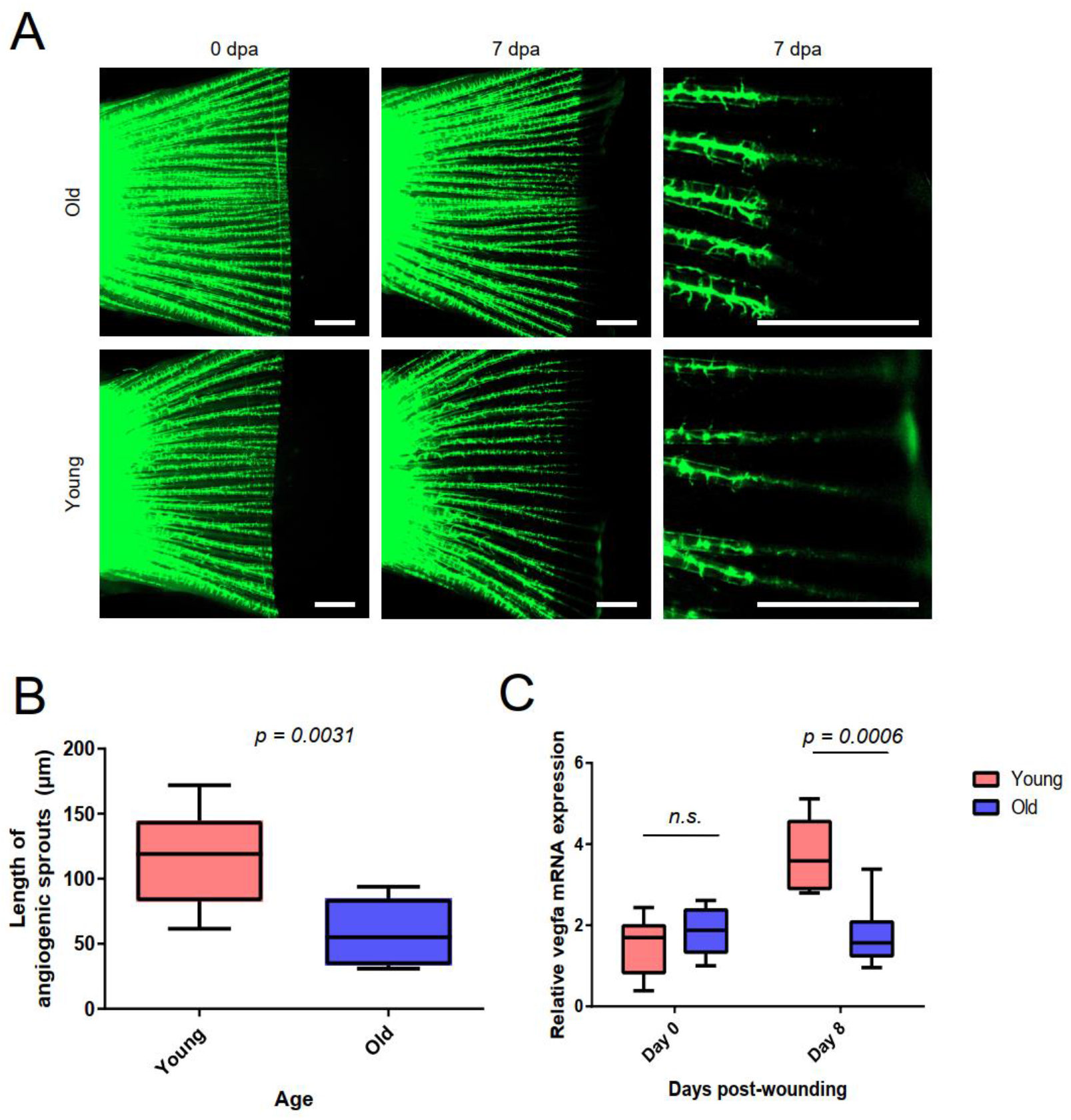
Angiogenesis is reduced in aged regenerating caudal fins. Zebrafish (genotype *roy, mitfa, Tg(fli1:EGFP)*) were subjected to caudal fin angiogenesis assay. Old fish were 36 weeks and young fish 16 weeks. A) Fluorescence microscopy images of transgenic zebrafish caudal fins during wound healing experiment. Statistical analysis with Mann-Whitney U-test. B) Quantification of angiogenesis in transgenic zebrafish caudal fins. Old fish (n=8), young fish (n=10). C) qRT-PCR analysis of *vegfa* mRNA expression during wound healing assay in turquoise killifish. n=6 in each group. Statistical analysis was done with two-way ANOVA, age and time used as factors and multiple comparison testing with Sidak’s post-hoc test.

To further characterize changes in the vasculature, we analysed the expression of key angiogenic growth factor *vegfa* using qPCR. The expression of *vegfa* was induced in young regenerating fins, whereas this response was missing in old killifish (Fig. 4C).

These results indicated that alterations in vasculature and angiogenic *vegfa* signalling may underlie the aging-related decline in wound healing and fin regeneration.

### Regeneration of aged caudal fin is inhibited by anti-angiogenic drug

These data indicated that dysregulated angiogenesis may play a role in reduced regeneration of caudal fin in aged individuals. In young adult zebrafish, the fin regeneration is dependent on angiogenesis and pharmacological inhibition of VEGF signalling can reduce fin regeneration ^27^. To address if VEGF signalling is still required in fin regeneration of aged fish, we treated the killifish with VEGF-receptor inhibitor SU5416 ^28^ after resection of the caudal fin. Indeed, this treatment reduced the growth of caudal fins in aged killifish (Fig 5A,B,C). The observation of effectiveness of SU5416 to alter fin regeneration in aged killifish is in-line with the earlier observations in young adult zebrafish ^27^. As expected, we observed also reduced number of perfused vessels in histological sections (Fig. 5D). To confirm this further, we analysed vascular biomarkers *vegfa* and *cd34* using qPCR. The reduced expression of vascular biomarkers *cd34* (Fig. 5E) and *vegfa* (Fig. 5F) indicated the effectiveness of anti-angiogenic SU5416 treatment in targeting vasculature.

**Figure 5.**
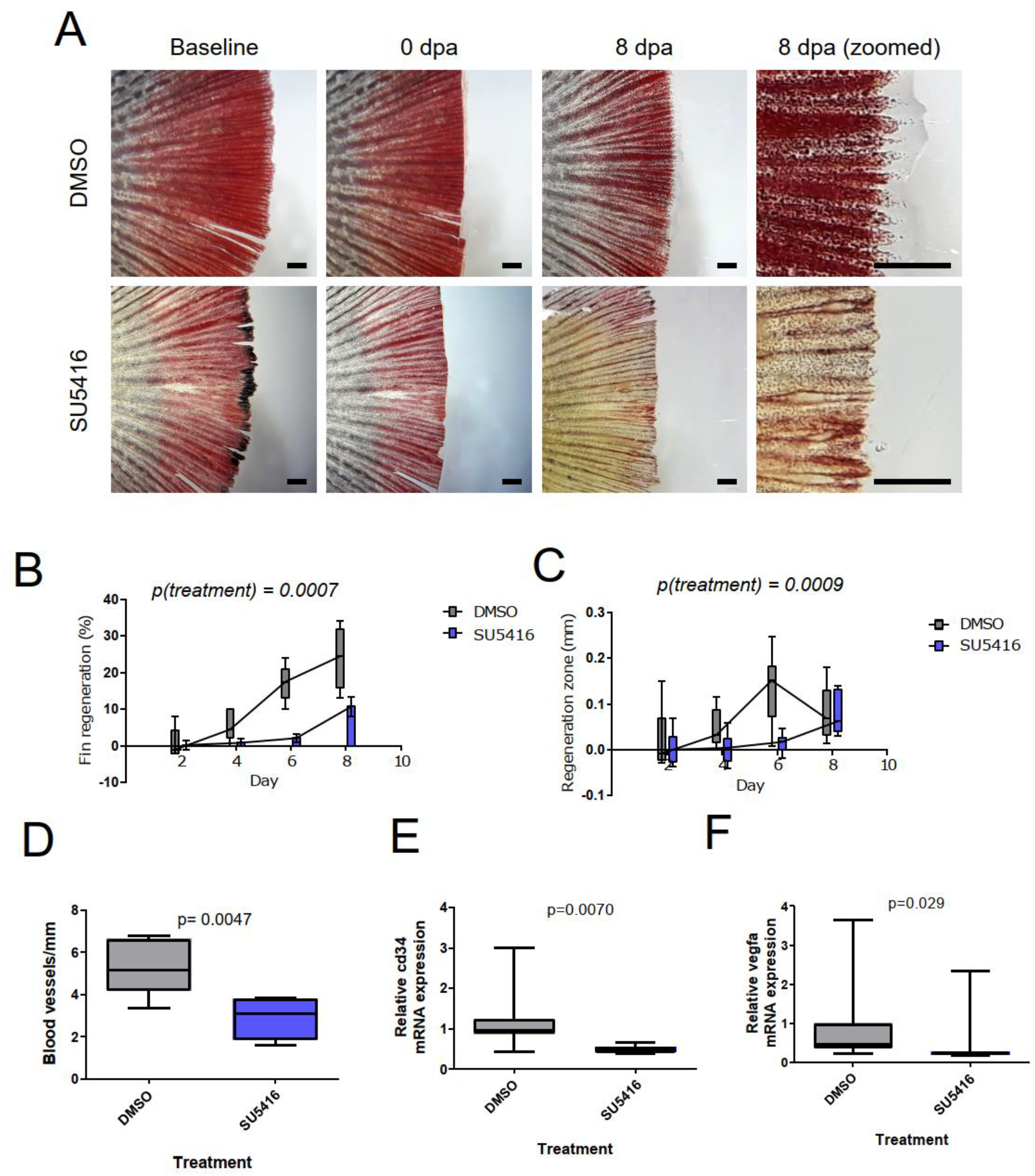
Anti-angiogenic drug SU5416 reduces caudal fin regeneration in aged fish. Caudal fin angiogenesis assay with old turquoise fish and control or SU5416 treatment. A) Microscopy images of caudal fins of turquoise killifish during caudal fin regeneration assay n=6 for DMSO and n= 8 for SU5416. Scale bar 1mm. B) Quantification of total regeneration of caudal fin. C) Quantification of thickness of regenerating area. D) Measurement of blood vessel density in histological sections. E, F) qPCR measurements of mRNA expression of vascular biomarkers *cd34* (E) and *vegfa* (F). n=7 for both groups. Statistical analysis in C and D is 2-way ANOVA, and in D, E, F Mann-Whitney U-test.

### SU5416 alters vascular biomarkers, collagen deposition and cellular proliferation

To gain more in-depth analysis of tissue regeneration, we carried out immunostainings of regenerating fins (Fig. 6A). Interestingly, the amount of collagen was reduced (Fig. 6B) indicating defects in the regeneration of the caudal fin tissue structures. Abnormally high cell proliferation rates were observed after SU5416 treatment (Figs 6C), and these proliferating cells were localized throughout the fin tissue. This indicates that the effects of the inhibition of VEGF signalling extend beyond vascular signalling and angiogenesis in regenerating fins.

**Figure 6.**
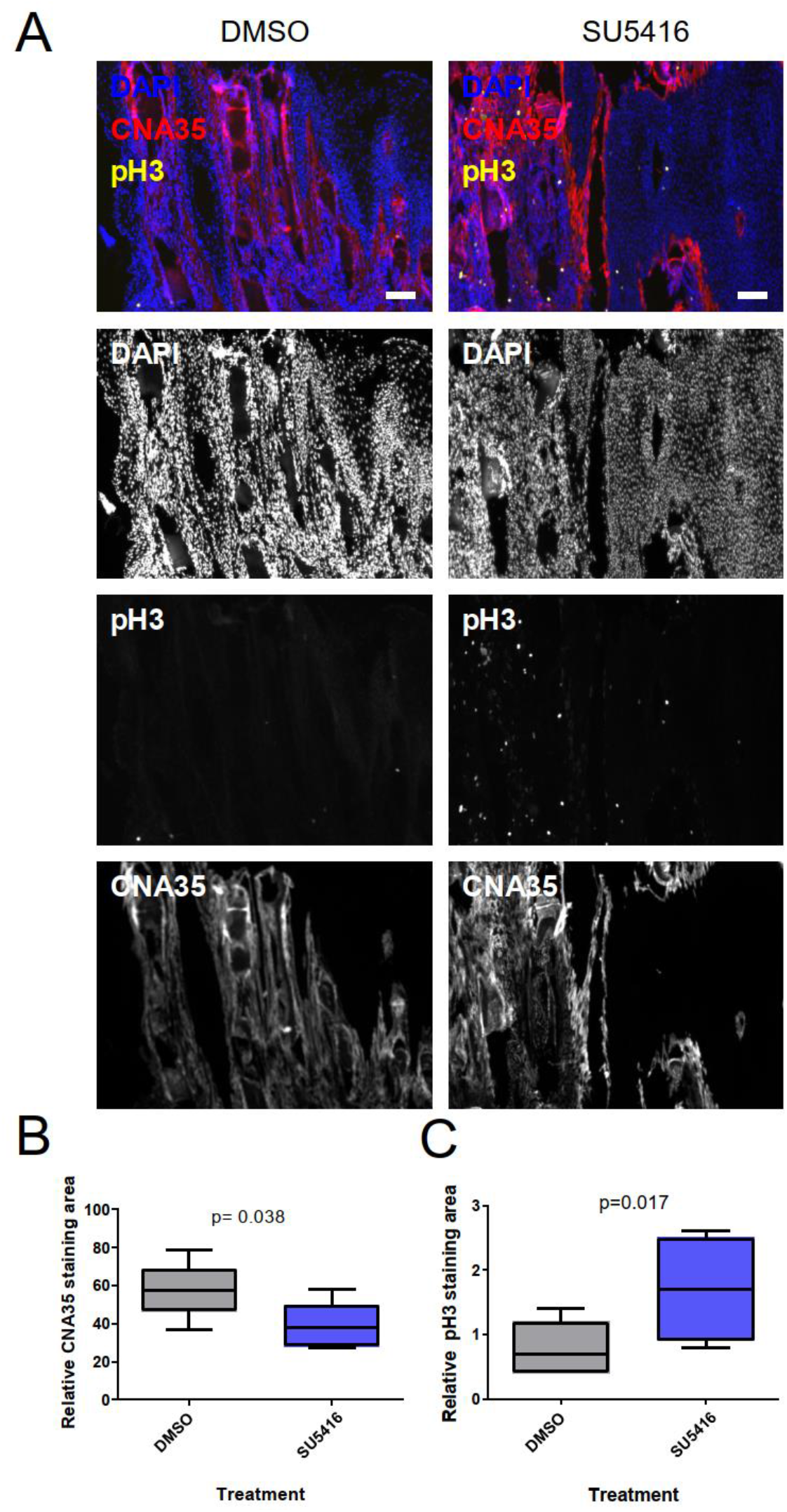
Anti-angiogenic drug SU5416 alters cellular response during caudal fin regeneration. Samples were obtained from caudal fin regeneration assay and processed to histological sections. A) Image of immunostaining for proliferation (pH3), collagen (CNA35) and nuclei (DAPI). Scale bar 0.1mm. B) Quantification of collagen content stained by CNA-35 probe. C) Quantification of cell proliferation marker phospho-histone 3 (pH3). n=7 in both groups.

## Discussion

In our experiments, we observed that aging results in declined wound healing and fin regeneration in both zebrafish and turquoise killifish. This is in line with the earlier observations with wound healing model of turquoise killifish ^24^. The previous results in zebrafish have been more controversial, and both decline and normal wound healing have been observed ^29,30^. One explanation could be in the long lifespan of zebrafish. Zebrafish can live up to 5 years in laboratory, but 2-year old fish are often considered middle-aged to old and begin to show signs of aging in aging studies ^31^. Although aging is a gradual process and aging-related changes occur well-before reaching maximal lifespan, the pace of aging varies across phenotypic traits ^32^. Hence, using fish much closer to the actual maximal life span could provide clearer results. This, however, causes problems in carrying out the experiments due to long aging period before obtaining suitable populations.

Turquoise killifish is much more amenable to aging studies due to its shorter natural lifespan ^17^. Fish strain MZCS-222 used in this study had median lifespan of 34 weeks when housed in Turku, Finland. It is also important to note that many factors in husbandry may affect lifespan as conditions in different laboratories may differ in numerous ways ^19^. The same strain housed in Brno, Czech Republic had lifespan of 18-25 weeks. The factors underlying the variation involve temperature, diet, water quality parameters, frequency of mating and mycobacterial infection ^19,33^. Therefore, it is important to understand the aging dynamics also in the laboratory carrying out the experiments.

During these experiments, we observed increased vascular density and controversially also a reduced angiogenic potential in aged fish. The increased vascular density was likely a secondary effect, as the vessels were tightly associated with fin rays which were more branched in aged fish. The reduced angiogenesis was, however, functionally significant. The blockage of VEGF signalling had significant effect on wound healing in aged turquoise killifish, indicating that despite reduced angiogenesis and VEGF expression, the VEGF signalling system remains functional and physiologically relevant also in the caudal fins of aging fish. This observation is supported by earlier results of impaired VEGF signalling during aging process in mammals^14^. Our findings imply a central role for angiogenesis during various processes of fin regeneration extending beyond the growth of new blood vessels. Many of these effects are likely to arise secondarily due the lack of functional vasculature to support new regenerating tissues with sufficient oxygen and nutrients.

Interestingly, poor angiogenesis has been implicated also with poor wound healing in humans ^34^. This underlines the need to obtain deeper mechanistic understanding of the mechanisms of both angiogenesis and wound healing in aged tissues. The aging-related changes in angiogenesis are still inadequately characterized and there is a lack of suitable models to characterize them ^15^. Our data indicates that the wound healing models of turquoise killifish and zebrafish may be suitable models to characterize the role of and mechanisms of angiogenesis in wound healing of aged animals. This can pave the way for future identification of suitable angiogenic agents with potential to enhance wound healing in aged individuals.

## Acknowledgements

We thank Cell Imaging Core, Zebrafish Core and Finnish Functional Genomics Centre (Turku Bioscience Centre, University of Turku and Åbo Akademi University, Turku, Finland) supported by Biocenter Finland for expertise. We thank HistoCore (University of Turku) for processing samples for histology and H&E staining. Image data management was supported by the Turku BioImaging Image Data Team at Åbo Akademi University and University of Turku.

## Funding

Turku University Foundation (I.P.).

## Competing Interests

The authors have no relevant financial or non-financial interests to disclose.

## Author Contributions

All authors contributed to the study conception and design. Material preparation, data collection and analysis were performed by J.Ö., K.K., J.V. and I.P. Killifish lines were selected and provided by M.R. As corresponding author, I.P. had full access to all the data in the study and takes responsibility for its integrity and the data analysis. The first draft of the manuscript was written by J.Ö. and I.P., and all authors commented the manuscript. All authors read and approved the final manuscript.

## Data Availability

Data is available from authors upon reasonable request.

## Ethics approval

The authorization for this project was obtained from Animal Experimental Board of Regional State Administrative Agency for Southern Finland (ESAVI/16458/2019 and ESAVI/2402/2021). The experimentation was carried out following Directive 2010/63/EU and associated Finnish legislation.

## Highlights

- **Zebrafish and turquoise killifish are complementing models for aging studies.**
- **Aging reduces angiogenesis in regenerating caudal fin.**
- **Anti-angiogenic treatment reduces regeneration of aged caudal fin.**

## Notes

### Competing Interest Statement

The authors have declared no competing interest.

## References

1. Sgonc R, Gruber J. Age-related aspects of cutaneous wound healing: A mini-review. Gerontology. 2013;59(2):159–164. doi:10.1159/000342344

2. Sen CK, Gordillo GM, Roy S, et al. Human skin wounds: A major and snowballing threat to public health and the economy: PERSPECTIVE ARTICLE. Wound Repair Regen. 2009;17(6):763–771. doi:10.1111/j.1524-475X.2009.00543.x

3. Posnett J, Franks PJ. The burden of chronic wounds in the UK. Nurs Times. 2008;104(3):44–45.

4. Phillips CJ, Humphreys I, Fletcher J, Harding K, Chamberlain G, Macey S. Estimating the costs associated with the management of patients with chronic wounds using linked routine data. Int Wound J. 2016;13(6):1193–1197. doi:10.1111/iwj.12443

5. Frykberg RG, Banks J. Challenges in the Treatment of Chronic Wounds. Adv Wound Care. 2015;4(9):560–582. doi:10.1089/wound.2015.0635

6. Honnegowda TM, Kumar P, Padmanabha Udupa EG, et al. Effects of limited access dressing in chronic wounds: A biochemical and histological study. Indian J Plast Surg. 2015;48(1):22–28. doi:10.4103/0970-0358.155263

7. Hellberg C, Ostman A, Heldin C-H. PDGF and vessel maturation. Recent results cancer Res Fortschritte der Krebsforsch Prog dans les Rech sur le cancer. 2010;180:103–114. doi:10.1007/978-3-540-78281-0_7

8. Li X, Eriksson U. Novel VEGF family members: VEGF-B, VEGF-C and VEGF-D. Int J Biochem Cell Biol. 2001;33(4):421–426. doi:10.1016/S1357-2725(01)00027-9

9. Sammarco MC, Dawson LA, Simkin J, Yu L, Muneoka K, Schanes PP. The mammalian blastema: regeneration at our fingertips. Regeneration. 2015;2(3):93–105. doi:10.1002/reg2.36

10. Acker T, Plate KH. Role of hypoxia in tumor angiogenesis - Molecular and cellular angiogenic crosstalk. Cell Tissue Res. 2003;314(1):145–155. doi:10.1007/s00441-003-0763-8

11. Kumar I, Staton CA, Cross SS, Reed MWR, Brown NJ. Angiogenesis, vascular endothelial growth factor and its receptors in human surgical wounds. Published online 2009:1484–1491. doi:10.1002/bjs.6778

12. Rivard A, Berthou-Soulie L, Principe N, et al. Age-dependent defect in vascular endothelial growth factor expression is associated with reduced hypoxia-inducible factor 1 activity. J Biol Chem. 2000;275(38):29643–29647. doi:10.1074/jbc.M001029200

13. Ashcroft GS, Mills SJ, Ashworth JJ. Ageing and wound healing. Biogerontology. 2002;3(6):337–345. doi:10.1023/a:1021399228395

14. Grunewald M, Kumar S, Sharife H, et al. Counteracting age-related VEGF signaling insufficiency promotes healthy aging and extends life span. Science (80-). 2021;8479. doi:10.1126/science.abc8479

15. Hodges NA, Suarez-Martinez AD, Murfee WL. Understanding angiogenesis during aging: Opportunities for discoveries and new models. J Appl Physiol. 2018;125(6):1843–1850. doi:10.1152/japplphysiol.00112.2018

16. Goswami AG, Basu S, Shukla VK. Wound Healing in the Golden Agers: What We Know and the Possible Way Ahead. Int J Low Extrem Wounds. Published online 2021. doi:10.1177/15347346211037841

17. Harel I, Brunet A. The African Turquoise Killifish: A Model for Exploring Vertebrate Aging and Diseases in the Fast Lane. Cold Spring Harb Symp Quant Biol. 2015;LXXX:275–279. doi:10.1101/sqb.2015.80.027524

18. Hu CK, Brunet A. The African turquoise killifish: A research organism to study vertebrate aging and diapause. Aging Cell. 2018;17(3):1–15. doi:10.1111/acel.12757

19. Polačik M, Blažek R, Reichard M. Laboratory breeding of the short-lived annual killifish Nothobranchius furzeri. Nat Protoc. 2016;11(8):1396–1413. doi:10.1038/nprot.2016.080

20. Aper SJA, Van Spreeuwel ACC, Van Turnhout MC, et al. Colorful protein-based fluorescent probes for collagen imaging. PLoS One. 2014;9(12):1–21. doi:10.1371/journal.pone.0114983

21. Allan C, Burel J-M, Moore J, et al. OMERO: flexible, model-driven data management for experimental biology. Nat Methods. 2012;9(3):245–253. doi:10.1038/nmeth.1896

22. Blažek R, Polačik M, Reichard M. Rapid growth, early maturation and short generation time in African annual fishes. Evodevo. 2013;4(1):24. doi:10.1186/2041-9139-4-24

23. Gerhard GS, Kauffman EJ, Wang X, et al. Life spans and senescent phenotypes in two strains of Zebrafish (Danio rerio). Exp Gerontol. 2002;37(8-9):1055–1068. doi:10.1016/S0531-5565(02)00088-8

24. Wendler S, Hartmann N, Hoppe B, Englert C. Age-dependent decline in fin regenerative capacity in the short-lived fish Nothobranchius furzeri. Aging Cell. 2015;14(5):857–866. doi:10.1111/acel.12367

25. Nachtrab G, Czerwinski M, Poss KD. Sexually dimorphic fin regeneration in Zebrafish controlled by androgen/GSK3 signaling. Curr Biol. 2011;21(22):1912–1917. doi:10.1016/j.cub.2011.09.050

26. Hlushchuk R, Brönnimann D, Shokiche CC, et al. Zebrafish caudal fin angiogenesis assay-Advanced quantitative assessment including 3-way correlative microscopy. PLoS One. 2016;11(3). doi:10.1371/journal.pone.0149281

27. Bayliss PE, Bellavance KL, Whitehead GG, et al. Chemical modulation of receptor signaling inhibits regenerative angiogenesis in adult zebrafish. Nat Chem Biol. 2006;2(5):265–273. doi:10.1038/nchembio778

28. Fong TA, Shawver LK, Sun L, et al. SU5416 is a potent and selective inhibitor of the vascular endothelial growth factor receptor (Flk-1/KDR) that inhibits tyrosine kinase catalysis, tumor vascularization, and growth of multiple tumor types. Cancer Res. 1999;59(1):99–106. http://www.ncbi.nlm.nih.gov/pubmed/9892193

29. Tsai SB, Tucci V, Uchiyama J, et al. Differential effects of genotoxic stress on both concurrent body growth and gradual senescence in the adult zebrafish. Aging Cell. 2007;6(2):209–224. doi:10.1111/j.1474-9726.2007.00278.x

30. Itou J, Kawakami H, Burgoyne T, Kawakami Y. Life-long preservation of the regenerative capacity in the fin and heart in zebrafish. Biol Open. 2012;1(8):739–746. doi:10.1242/bio.20121057

31. Kishi S, Slack BE, Uchiyama J, Zhdanova I V. Zebrafish as a genetic model in biological and behavioral gerontology: Where development meets aging in vertebrates - A mini-review. Gerontology. 2009;55(4):430–441. doi:10.1159/000228892

32. Hayward AD, Moorad J, Regan CE, et al. Asynchrony of senescence among phenotypic traits in a wild mammal population. Exp Gerontol. 2015;71:56–68. doi:10.1016/j.exger.2015.08.003

33. Reichard M, Blažek R, Dyková I, Žák J, Polačik M. Challenges in keeping annual killifish. In: D’Angelo L., P de G, eds. Laboratory Fish in Biomedical Research. Academic Press; 2022:289–310. doi:10.1016/b978-0-12-821099-4.00001-8

34. Cheng B, Fu X. The Focus and Target: Angiogenesis in Refractory Wound Healing. Int J Low Extrem Wounds. 2018;17(4):301–303. doi:10.1177/1534734618813229

